# Genetic perturbation of mitochondrial function reveals functional role for specific mitonuclear genes, metabolites and pathways that regulate lifespan

**DOI:** 10.1101/2023.01.26.525799

**Authors:** Cheryl Zi Jin Phua, Xiaqing Zhao, Lesly Turcios-Hernandez, Morrigan McKernan, Morteza Abyadeh, Siming Ma, Daniel Promislow, Matt Kaeberlein, Alaattin Kaya

## Abstract

Altered mitochondrial function is tightly linked to lifespan regulation, but underlying mechanisms remain unclear. Here, we report the chronological and replicative lifespan variation across 168 yeast knock-out strains, each lacking a single nuclear-coded mitochondrial gene, including 144 genes with human homologs, many associated with diseases. We dissected the signatures of observed lifespan differences by analyzing profiles of each strain’s proteome, lipidome, and metabolome under fermentative and respiratory culture conditions, which correspond to the metabolic states of replicative and chronologically aging cells, respectively. Examination of the relationships among extended longevity phenotypes, protein, and metabolite levels revealed that although many of these nuclear-encoded mitochondrial genes carry out different functions, their inhibition attenuates a common mechanism that controls cytosolic ribosomal protein abundance, actin dynamics, and proteasome function to regulate lifespan. The principles of lifespan control learned through this work may be applicable to the regulation of lifespan in more complex organisms, since many aspects of mitochondrial function are highly conserved among eukaryotes.

## Introduction

Mitochondria are semi-autonomous organelles that utilize glucose and fatty acids through oxidative phosphorylation to generate energy (in the form of ATP) for eukaryotic cells. In addition, mitochondria produce key metabolites and biochemical signals, regulate cellular homeostasis, and carry out essential processes such as apoptosis and autophagy [1–3]. With rare exceptions [4], eukaryotic cells without mitochondria are not viable [5].

Although several genes encoded by the mitochondrial genome (mtDNA) regulate different steps in oxidative phosphorylation and cellular metabolic regulation (~1% of mitochondrial genes), mitochondrial biogenesis is also dependent on 1,000-1,200 nuclear (nDNA) encoded proteins localized to the mitochondria [6–8]. Accordingly, defects in nDNA-encoded mitochondrial genes are associated with a multitude of mitochondrial dysfunctions, contributing to many diseases, including primary inherited mitochondrial disorders, Parkinson’s disease, diabetes mellitus, and Alzheimer’s disease [9–12]. In addition, aging is marked by a decrease in mitochondrial respiratory capacity, biogenesis, mtDNA copy number, and a decline in the activity of mitochondrial enzyme complexes. Accordingly, the decline in mitochondrial function is associated with many hallmarks of aging and age-related diseases [13–16]. Conversely, several genetic, environmental, and pharmacological interventions have been shown to be intertwined with mitochondria-related pathways to mediate longevity*-*promoting metabolic changes in different organisms, including yeast [17–19] *C. elegans* [20, 21], and mice [22, 23]. Intriguingly, inhibition of mitochondrial respiration has also been associated with increased lifespan in various species, including yeast [24], fly [25], worms [26], and mice [27]. For example, several worm strains with knockdown of genes encoding mitochondrial proteins exhibit a mild decrease in mitochondrial respiration and extended lifespan [28]. Overall, a growing body of data from studies on model systems demonstrates that the plasticity of mitochondrial function (mild decrease or increase in respiration) could be a potential target to promote healthy aging. However, genes and related regulatory networks causally linking these changes to longevity remain unclear. While there has been an explosion of research focusing on the genetic alteration of individual mitochondrial proteins in altering mitochondrial function, we are still missing the mechanisms underlying varying degrees of improvement or reduction in mitochondrial function in regulating aging.

As the organism in which mtDNA was first sequenced [29, 30], the budding yeast *Saccharomyces cerevisiae* (*S. cerevisiae*) has played an important role in understanding how mitochondria interact with various cellular pathways and functions. Most mitochondrial functions and nuclear-coded mitochondrial genes (~900 in yeast and ~1500 in human) are highly conserved across eukarya [31–33]. Moreover, being a facultative-anaerobe (producing energy primarily from fermentation), *S. cerevisiae* can survive with mutations that inactivate oxidative phosphorylation or tolerate the complete loss of mtDNA in the presence of a fermentative substrate [34]. As a result, modification of the mitochondrial genome or mitochondrial proteins are often studied without cell lethality. These specific characteristics of its metabolism and mitochondrial biology have allowed functional genomic studies of *S. cerevisiae* to identify mitochondrial genes involved in human diseases, and to uncover a central role for mitochondrial function in aging [32].

To better understand the mechanisms linking mitochondrial function with lifespan variation, we characterized how the loss of individual conserved nuclear-encoded mitochondrial proteins influenced cellular physiology and longevity. We analyzed both the chronological (CLS) and replicative life span (RLS) of 168 yeast knock-out strains, each lacking one mitochondrial protein (mtKO) [35]. We found a wide range of variation in both CLS and RLS and then dissected the signatures of the observed CLS and RLS differences by analyzing the correlation and interaction between protein, lipid, and metabolite abundance with the observed lifespan differences. This approach allowed us to uncover the sets of genes and metabolites that are associated with, and might have a causative role in, lifespan regulation. In addition, integrating different levels of “omics” data led to a systems-level view of the molecular changes in key pathways involved in longevity regulation. Our analyses indicated that modulation of cytosolic and mitochondrial ribosomal protein abundance, actin dynamics, and proteasome capacity of the cells, tightly regulated by the functional status of mitochondria, and the balance in these processes, affected cellular aging. In addition, we found that metabolic fates of glutamate through orotate and alphaketoglutarate might have a significant role in regulating lifespan.

## Results

### Lifespan variation across the 168 deletion strains each lacking a single conserved gene related to mitochondrial function

For each mitochondrial gene knockout (mtKO) strain, we characterized two forms of lifespan: chronological lifespan (CLS, i.e., survival time of populations of non-dividing cells) and replicative lifespan (RLS, i.e., number of daughter cells produced by a mother cell before senescence) (**Fig. 1A and Supplementary Table 1**). Across the 168 deletion strains, CLS ranged from 3 to 58 days, with ~70% of the strains living shorter than wild-type (WT, 41 days) (**Fig. 1A and Supplementary Table 1**). Among the shortest-lived were the three strains lacking mitochondrial ATP synthase (*Δatp12*: 3 days; *Δatp1*: 5 days; *Δatp2*: 19 days), those affecting mitochondrial electron transport chain complexes III/IV or their assembly (*Δafg3, Δcox12, Δqcr7, Δrip1, Δshy1*; 10 – 31 days), as well as eight out of the ten strains defective in biosynthesis of ubiquinone (*Δcoq1, Δcoq2, Δcoq3, Δcoq4, Δcoq6, Δcoq9, Δcoq11*; 5 – 27 days). These defects illustrate the crucial role of mitochondrial electron transport chain and cellular energetics in the regulation of CLS. The deletion strains affecting mitochondrial ribosomes or mitochondrial translation (*Δmrp1, Δmrp7, Δmrpl11, Δmef1, Δmef2*; 7 – 32 days) also tended to be short-lived. On the other hand, one of the longest-lived strains involved the deletion of a subunit of the TOR (Target of Rapamycin) complex 1 (*Δtor1*: 51 days), consistent with published literature that inhibition of the TOR signaling pathway is known to extend lifespan across multiple model organisms [36, 37], and deletion of TOR-pathway genes in yeast has been shown to significantly increase CLS previously [17, 38, 39]. Mic26p is a component of the mitochondrial inner membrane complex (MICOS) required for maintenance of the cristae junctions, and its deletion produced the longest CLS (*Δmic26*: 58 days). Disruption of mitofusin and its associated proteins also led to CLS extension effects (*Δfzo1*: 45 days; *Δmdm30*: 51 days). Energy generation during CLS of yeast (stationary phase culture) is fully dependent on mitochondrial function, since ethanol is the primary carbon source and requires functional mitochondria [40]. Accordingly, respiratory deficient mtKO strains were characterized by reduced CLS (**Fig. 1A and Supplementary Table 1**).

**Figure 1:**
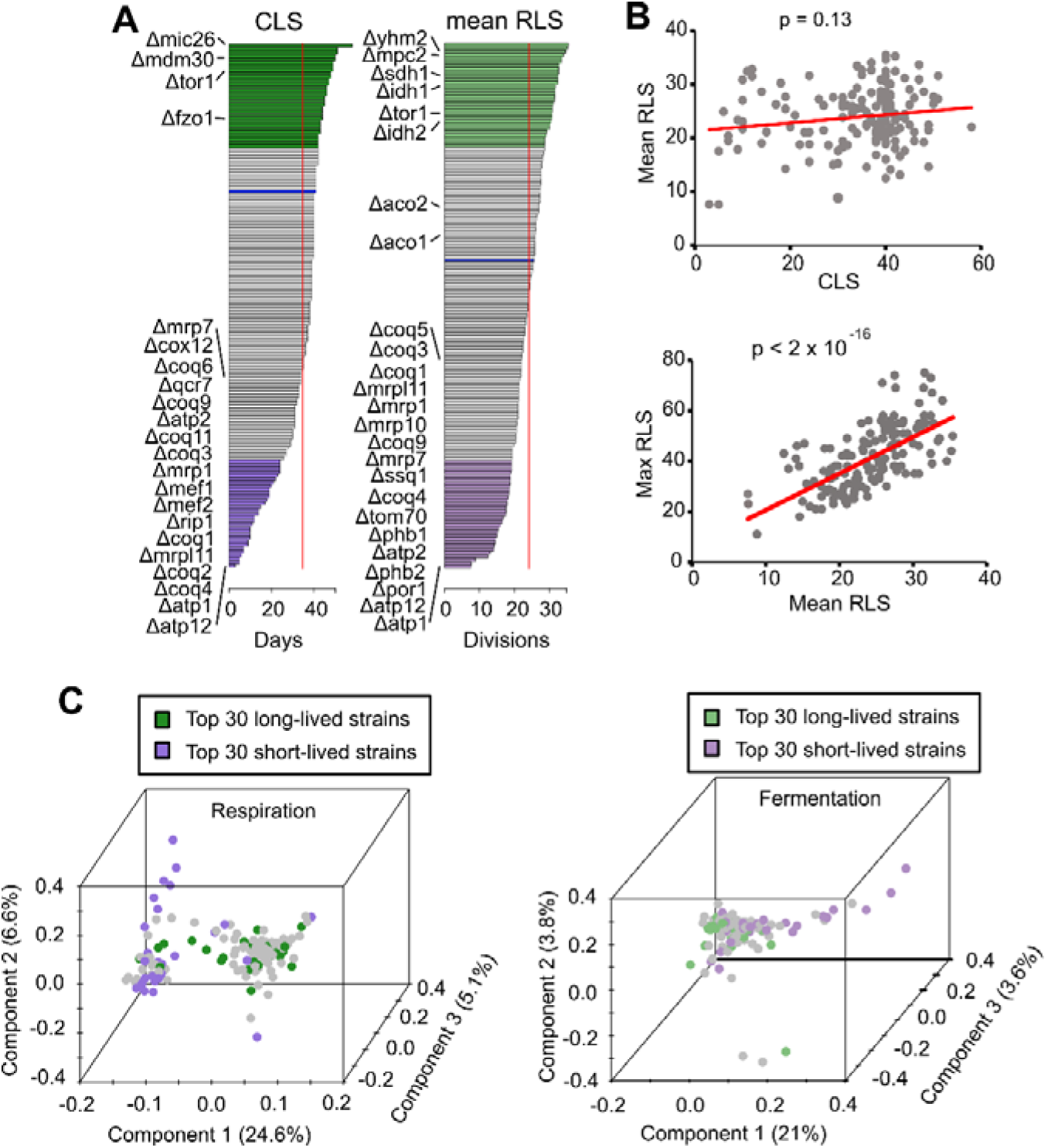
Phenotypic variation across mtKO strains. (**A**) CLS and mean RLS variation. Green and purple bars show top long- and short-lived strains, respectively. Blue bars indicate the lifespan of the WT control. Vertical red lines depict the average CLS (34 days) and mean RLS (24 divisions) across mtKO strains. (**B**) Correlation between different lifespan measurements. (**C**) Principal component analysis (PCA) of combined data. PCA was performed by combining proteins (peptides), metabolites and lipids data. The percentage variance explained by each principal component (PC) is shown in parentheses. The strains are colored using the same scheme as panel A. See Supplementary Figure 2 for separate PCA plots on each class of phenotype data.

Mean RLS ranged from 7.6 to 35.4 divisions, with ~56% of the strains living shorter than WT (25.2 divisions) (**Fig. 1A and Supplementary Table 1**) (Pearson correlation coefficient between mean and median RLS = 0.7, p < 2×10-16, Fig. 1B). Similar to CLS, those strains with mitochondrial ATP synthase deletions also had the shortest mean RLS (Δ*atp12* and *Δatp1*: 7.6 divisions; *Δatp2*: 14.2 divisions), which is consistent with prior reports, and suggests that maintenance of mitochondrial membrane potential during ATP production might be an important regulator of cellular aging [41–43]. Further underlying the key role of mitochondria in lifespan control, strains with a deletion in mitochondrial ribosomal proteins (Δ*mrp1, Δmrp7, Δmrp10, Δmrpl11*; 18.6 – 19.4 divisions) or ubiquinone biosynthesis (*Δcoq1, Δcoq3, Δcoq4, Δcoq5, Δcoq9*; 17.5 – 21.7 divisions) were generally short-lived, though the effects were not as large. Deletions of mitochondrial inner membrane chaperone prohibitins (*Δphb1*: 14.5 divisions; *Δphb2*: 14.1 divisions) led to a significant reduction in RLS as previously reported [44, 45]; so, too, did the disruption of the mitochondrial Hsp70-type chaperone (*Δssq1*: 17.9 divisions) and mitochondrial outer membrane proteins such as Por1 and Tom70 (*Δpor1*: 12.4 divisions; *Δtom70*: 16.6 divisions) (**Supplementary Table 1**). On the other hand, disruption of *TOR1* and *IDH2* (coding for isocitrate dehydrogenase subunit 2) were previously shown to improve RLS [46], and these deletion strains were indeed among the longest-lived (*Δtor1*: 31.6 divisions; *Δidh2*: 30.8 divisions). Deletion of other tricarboxylic acid (TCA) cycle enzymes, including succinate dehydrogenase (*Δsdh1*: 32.8 divisions), isocitrate dehydrogenase (*Δidh1*: 32.6 divisions), and aconitase (*Δaco1*: 26.2 divisions; *Δaco2*: 27.4 divisions), led to varying degrees of RLS extension (**Supplementary Table 1**).

Overall, our analysis of CLS and RLS characteristics of 168 mtKOs revealed that deletion of only three genes (*IDH2, IMO32*, and *TOR1*) caused significant extension in both CLS and RLS compared to the WT strain. In fact, we did not observed significant correlation (p= 0.001) between CLS and mean RLS of the mtKO strains (**Fig. 1B**). In fact, the CLS and RLS exhibit different molecular trajectories that the preferences between respirations versus glycolysis and the culture condition differences can affect distinct cellular processes and that components (genes) of these pathways might be differentially regulated under each aging model.

### Molecular patterns correlating with CLS

CLS in yeast can be defined as the period of survival in the stationary growth phase in which the non-dividing yeast cell population remains viable after glucose exhaustion [38, 40]. In this culture condition, the cells depend more heavily on respiration for ATP production, and survival is influenced by fermentative production of organic acids during the initial fermentative phase [47]. Accordingly, we aimed to explain the molecular signatures of CLS differences among these mtKO strains by analyzing their genome-wide differences in protein (proteomics), metabolite (metabolomics), and lipid (lipidomics) abundances identified under respiratory conditions (see Materials and Methods) [35]. We performed a Principal Component (PC) analysis on the combined data set (**Fig. 1C**) as well as each data type (**Supplementary Fig. 1**) to visualize phenotypic variations among the mtKO strains. PC analysis of the combined data set revealed a characteristic pattern, with the first three PCs explaining ~37 % of the total variance (**Fig. 1C**). PC analysis on the individual omics data (proteome, metabolome, and lipidome) also showed similar structures with the first three PC explaining ~42-57% of the total variance. Interestingly, along PC1, the strains are mainly segregated according to their CLS phenotypes in all three data types (**Fig. 1C and Supplementary Fig. 1**). When we segregated the deletion strains and proteins by hierarchical clustering, the deletion strains also formed two clusters with significant differences in lifespan (p = 7.65×10^-13^), which resembled the structure observed in PC analysis (**Fig. 2B**). In particular, the short-lived group (n=53, which included 25 out of the top 30 shortest-lived strains) consisted predominantly of strains impaired in mitochondrial ATP synthase (*Δatp1, Δatp2, Δatp12*), electron transport chain complexes (*Δafg3, Δcox12, Δqcr7, Δrip1, Δshy1*), mitochondrial ribosomal proteins (*Δmrp1, Δmrp7, Δmrp10, Δmrpl11*) and elongation factors (*Δmef1, Δmef2*), as well as ubiquinone biosynthesis (*Δcoq1, Δcoq2, Δcoq3, Δcoq4, Δcoq5, Δcoq6, Δcoq7, Δcoq9, Δcoq10*) (**Fig. 1A**), which were consistent with the changes in protein levels (**Fig. 3**). In other words, under respiratory conditions, the deletion of these mitochondrial genes led to a drastic reduction in oxidative phosphorylation and ATP generation, severe disruption of mitochondrial protein translation, and suppression of the TCA cycle and metabolite biosynthesis in short-living strains.

**Figure 2:**
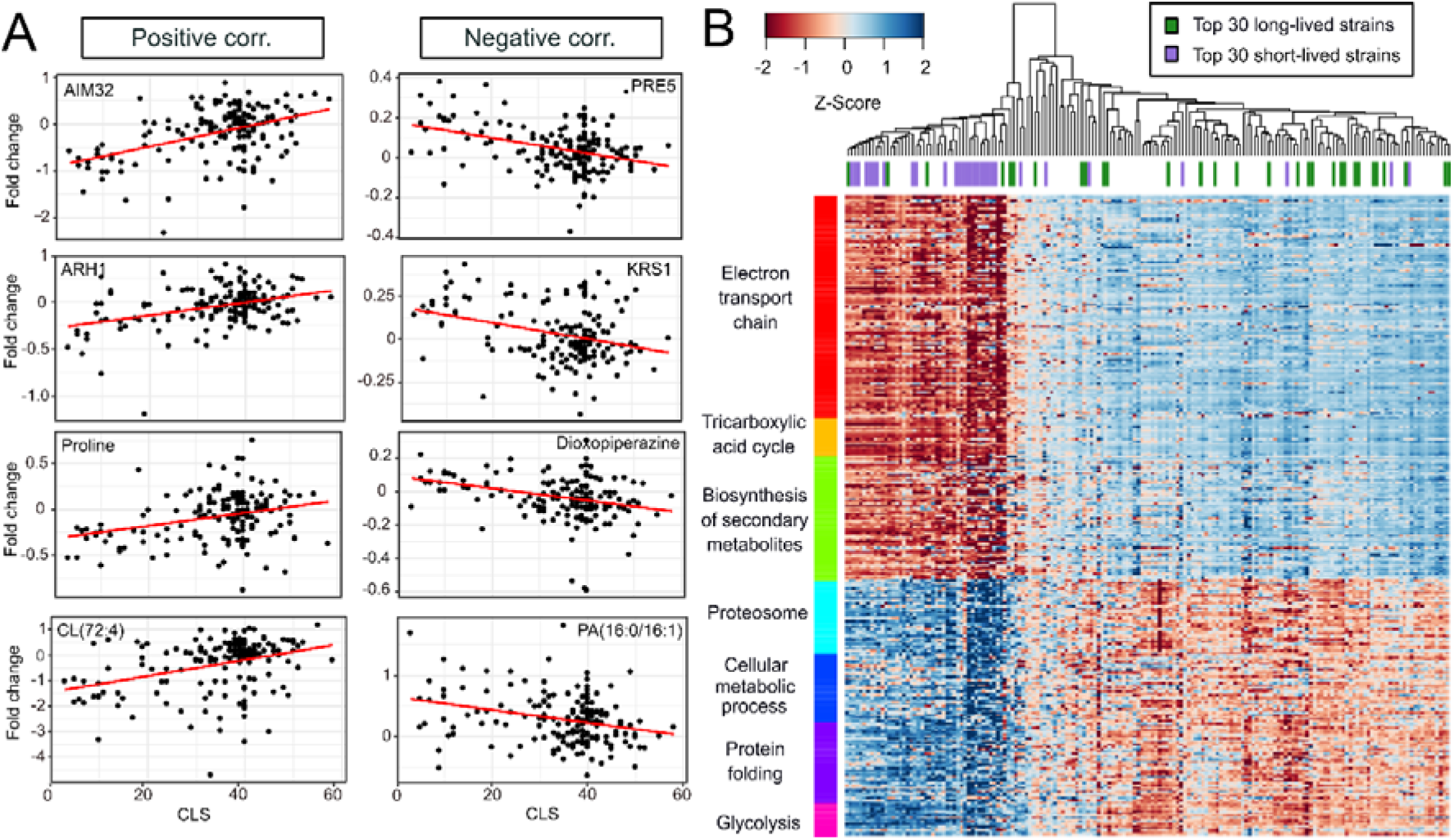
Selected endo phenotypes correlating with CLS. (**A**) Examples of proteins, metabolites, and lipids showing significant positive (left panel) or negative (right panel) correlation to CLS. P values and raw data can be found in Supplementary File 2. (**B**) The heat map on the right panel shows expression patterns of the significantly correlating proteins and enriched pathways. Spearman correlation matrix of aggregated protein expression profiles across strains was used for clustering. The top 30 short- and long-lived strains are indicated with purple and green colors (top panel on the heat map) as in Figure 1.

**Figure 3:**
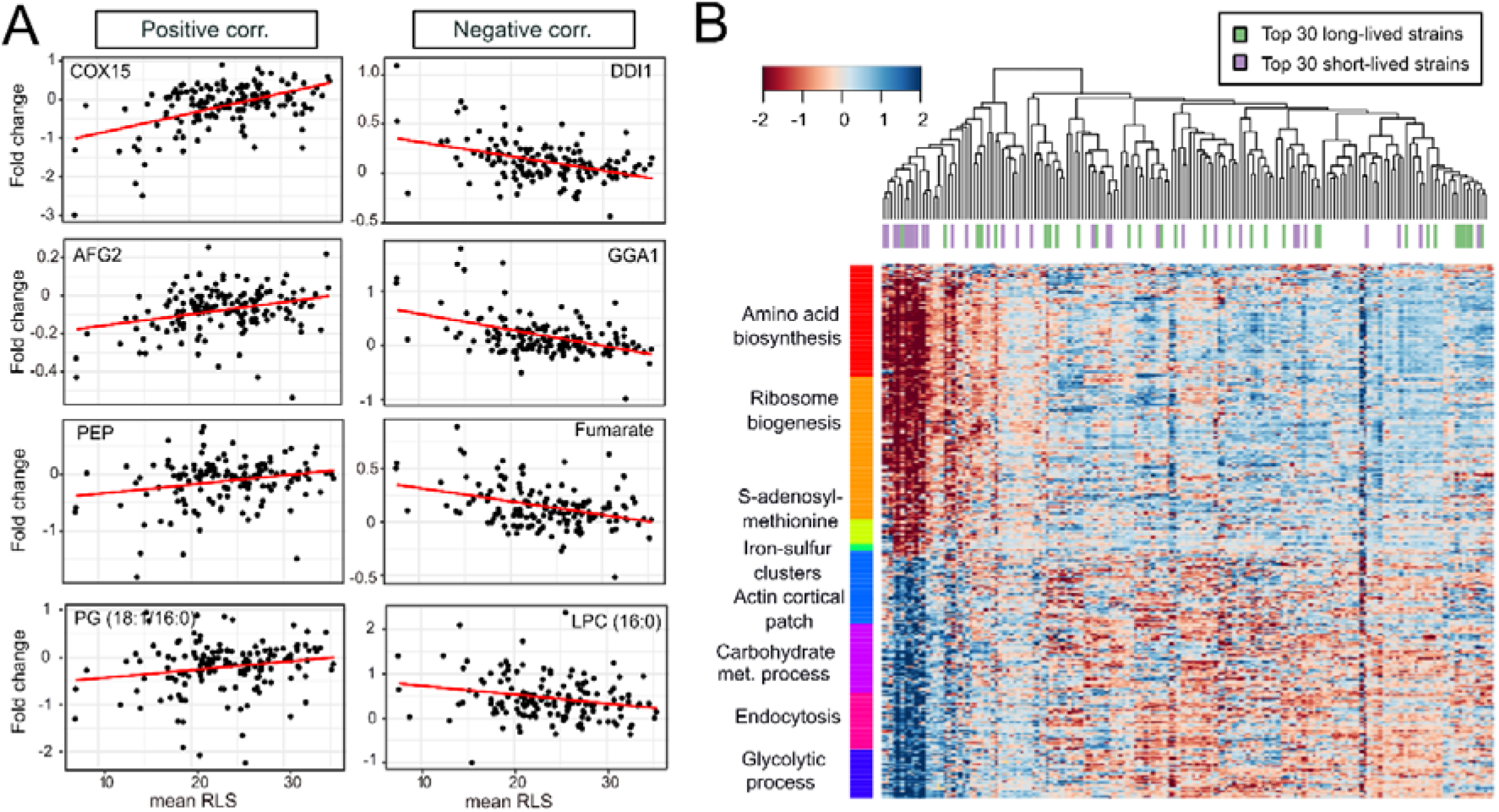
Selected endo phenotypes correlating with mean RLS. (**A**) Examples of proteins, metabolites, and lipids showing significant positive (left panel) or negative (right panel) correlation to mean RLS. P values and raw data can be found in Supplementary File 2. (**B**) The heat map on the right panel shows expression patterns of the significantly correlating proteins and enriched pathways. Spearman correlation matrix of aggregated protein expression profiles across strains was used for clustering. The top 30 short- and long-lived strains are indicated with purple and green colors (top panel on the heat map) as in Figure 1.

To further understand the molecular patterns underlying the CLS variations, we performed linear regression to identify proteins, metabolites, and lipids with significant correlation to the CLS of the mtKO strains. We identified 647 molecules (567 proteins, 59 metabolites, and 21 lipids) significantly correlated with CLS. (positive correlation: 313; negative correlation: 334; padj ≤ 0.01) (**Supplementary Table 2**). To address any potential imbalance in the numbers of short-lived strains and long-lived strains, we also repeated the analysis by using only the 30 shortest-lived (all with CLS shorter than wild-type) and 30 longest-lived (all with CLS longer than wild-type) deletion strains and found that 75% of the top molecules still remained significant at false discovery rate (FDR) < 0.01. Among the top proteins showing positive correlation, Arh1 (an oxidoreductase of the mitochondrial inner membrane; involved in cytoplasmic and mitochondrial iron homeostasis and required for activity of Fe-S cluster-containing enzymes), Ccp1 (a mitochondrial cytochrome-c peroxidase; degrades reactive oxygen species in mitochondria), and Aim32 (a 2Fe-2S mitochondrial protein involved in redox quality control) showed a positive correlation (**Fig. 2A and Supplementary Table 2**). Those showing negative correlation with CLS include: Dbp2 (an ATP-dependent RNA helicase of the DEAD-box protein family with transcription termination functions), Pre5 (alpha 6 subunit of the 20S proteasome), Krs1 (lysyl-tRNA synthetase), and New1 (a ribosome biogenesis factor-playing role in prion formation) (**Fig. 2A and Supplementary Table 2**).

Interestingly, while only ~18% of the negatively correlated genes were mitochondrial (localized to mitochondria), this number increased to ~74% for the positively correlated genes (**Supplementary Table 2)**. The positive correlation of these genes supported the observed CLS pattern. For example, we found three nuclear encoded mitochondrial proteins with distinct functions (Fmp21, Fmp25, and Fmp40), the abundance of which positively correlated with CLS. In fact, our analysis further verified this observation that null mutants of these three genes were characterized by decreased CLS, strongly suggesting their activity was required for CLS regulation (**Supplementary Table S1**). A similar observation could be seen for other proteins, whose deletion showed a strong CLS inhibition effect, such as Oct1, Atp1 and Atp12, Mrpl11, Coq5, Mef2, Kgd2, and Grx5, for which their null mutant decreased CLS by 87.5%, 92.5%, 82.5%, 75%, 72.5%, 47.5%, and 42.5%, respectively. On the other hand, there are six proteins (Mic26, Imo32, Aco2, Aro10, Kgd1, and Afg1) that their CLS phenotype contradict this observation: although their transcript levels positively correlated with CLS, their null mutant showed extended CLS phenotype. It is possible that for some of these genes, both their deletion and overexpression may positively affect CLS.

Pathway analyses revealed that the proteins with positive correlation to CLS were significantly enriched for “mitochondrial translation,” “mitochondrial gene expression,” “aerobic respiration,” and “ATP metabolic process.” In contrast, the proteins with a negative correlation to CLS were enriched for “cellular metabolic process,” “proteasome,” and “glycolytic process,” and response to chemicals (**Fig. 2B and Supplementary Table 2**). These data suggest that healthy mitochondria are necessary for maintaining longer CLS. Other studies have shown that increased proteasome capacity increases lifespan across different model organisms, but our data associate long-lived mtKO strains with decreased proteasome capacity. Therefore, we hypothesize that these long-lived mtKO strains might exhibit less accumulation of damaged (carbonylated and/or oxidized) proteins, resulting in reduced need for proteasome activity, which in turn leads to reduced energy expenditure and longer lifespan.

Observed changes in metabolites and lipids were generally consistent with the proteome observations. The levels of glycolytic and TCA cycle intermediates, including 2- phosphoglyceric acid (2PG), 3-phosphoglyceric acid (3PG), phosphoenolpyruvate (PEP), alpha-ketoglutarate (α-KG), proline and orotic acid all showed a significant positive correlation with CLS, whereas the amounts of glyceric acid, 2-hydroxyglutarate, and dioxopiperazine and lactic acid showed negative correlations (**Fig. 2A and Supplementary Table 2**). In addition, several nucleotides and nucleosides negatively correlated with CLS, including guanine, guanosine, adenosine, and AMP. With glycerol as the primary carbon source, the WT strain typically phosphorylates glycerol to glyceric acid, which is then oxidized to dihydroxyacetone phosphate (DHAP) using NAD^+^ or FAD^+^ as a proton acceptor. Subsequently, it enters glycolysis (appearing in the forms of 3PG, 2PG, and PEP) and the TCA cycle (α-KG) to support anabolic reactions [48]. In addition, 2-hydroxyglutarate can enter the TCA cycle as α-KG by donating a proton to NADP+. However, the mitochondrial electron transport chain was disrupted in the short-lived deletion strains. The inability to regenerate proton acceptors led to the accumulation of glycerol 3-phosphate and 2- hydroxyglutarate and the depletion of 2PG, 3PG, PEP, and α-KG. The levels of orotic acid, glutamine, glutamate, and asparagine derived from α-KG catabolism were also higher in the long-lived strains (**Supplementary Table 2**). These data indicate the beneficial effect of α- KG in CLS. It should also be noted that the abundance of many amino acids, including alanine, proline, isoleucine, and histidine, negatively correlated with CLS, suggesting decreased amino acid metabolism in long-lived strains.

In terms of lipids, 5 of the 11 cardiolipin species detected here correlated positively with CLS (**Fig. 2A and Supplementary Table 2**). Cardiolipin is found almost exclusively in the inner mitochondrial membrane and is required for the optimal activity of Complex I [49]. Furthermore, it was previously shown that the content of mitochondrial cardiolipin decreased, but the level of oxidized cardiolipin increased during aging; and that the age-related loss of Complex I activity in brain mitochondria could be restored by adding cardiolipin [50]. On the other hand, three phosphatidylserine (PS) and, all four cytidine diphosphate diacylglycerol (CDP-DAG) species correlated negatively with CLS (**Fig. 2A and Supplementary Table 2)**.

Overall, the up-regulation of mitochondrial translation, oxidative phosphorylation, and TCA cycle functions, as well as the down-regulation of protein degradation and amino acid metabolism, tend to affect CLS among mtKO strains positively. In contrast, changes in the opposite direction would often lead to shorter CLS.

### Molecular patterns correlating with RLS

We followed a similar approach to identify the proteins, lipids, and metabolites with significant correlation to RLS under fermentation conditions. Our PC analysis did not reveal clear segregation of strains on any component based on RLS variation (**Fig. 1C and Supplementary Fig. 2**). Although the first three PCs explained ~32-53% of the total variation (across each data), there might be other features discriminating separation among these mtKO strains. Next, we applied linear regression between molecular abundances and mean RLS. A total of 337 molecules (326 proteins, 10 metabolites, and 1 lipid) were found to be significantly correlated with mean RLS (positive correlation: 202; negative correlation: 145; padj ≤ 0.01) (**Supplementary Table 2**). Similar to CLS, we repeated the analysis using only the molecules from the 30 most long-lived strains and the 30 most short-lived strains and found that 78% of the top molecules were still significant at FDR < 0.01.

We found that ~65% of the positively correlated proteins were nuclear proteins, particularly localized to the nucleolus, known as the site of ribosome biogenesis. Among the negatively correlating proteins, ~30% of them were cytoplasmic (cytoskeleton binding proteins), ~29% of them were mitochondrial, and ~19% of them were localized to the plasma membrane (**Supplementary Table 2**).

Among the top proteins that showed a significant positive correlation with mean RLS are: Cox15 (a heme a synthase), Utp21 (a protein involved in ribosomal small subunit processome complex), Afg2 (an ATPase involved in ribosome maturation and degradation of aberrant mRNAs), and Sda1 (a protein required for actin organization) (**Fig. 3A and Supplementary Table 2**). On the other hand, the following proteins are among the top hits showing significant negative correlation with mean RLS: Tps2 (a phosphatase complex protein involved in the synthesis of storage carbohydrate trehalose), Gga1 (a golgi-localized protein with homology to gamma-adaptin), Dcs1 (a hydrolase involved in mRNA decapping), and Ddi1 (an aspartic-type endopeptidase involved in proteasome-independent, ubiquitin-dependent proteolysis) (**Fig. 3A and Supplementary Table 2)**.

Pathway enrichment analysis revealed significant enrichment for terms such as “ribosome biogenesis,” “amino acid biosynthesis,” and iron-sulfur (Fe-S) clusters,” (**Fig. 3B and Supplementary Table 2**) for the proteins that are positively correlated with mean RLS. The proteins with significant negative correlation to RLS were enriched for “carbohydrate metabolic process,” “actin organization,” “glycolytic process,” and “endosome” (**Fig 3B and Supplementary Table 2**).

Half of the metabolites correlating with mean RLS consisted of a group of unidentified metabolites (5 out of 10). Among the identified metabolite species, only orotic acid (OA) showed a significant positive correlation. In contrast, metabolites negatively correlated with RLS consisted of adenine, malic acid, fumaric acid, and citric acid (**Fig. 3A and Supplementary Table 2),** which are downstream intermediates of the TCA cycle, indicating a decrease of TCA cycle activity in long-lived strains. This observation is also consistent with gene expression pattern and pathway analysis in which we found decreased TCA cycle activity in long-lived strains. Our analysis did not reveal any lipid species that showed a significant positive correlation with mean RLS. Phosphatidylglycerol (PG) was the only lipid species that negatively correlated with RLS (**Fig. 3A and Supplementary Table 2)**. Overall, our data suggest maintenance of cytosolic translation, mitochondrial respiration, Fe-S cluster biogenesis, and actin dynamics as an important regulator of RLS process in strains with altered mitochondrial function.

### Experimental testing of orotic acid and alpha ketoglutarate effects on lifespan

Our –omics data suggested that an increased abundance of orotic acid (OA) positively correlates with both CLS and RLS (**Fig. 4A**). Another metabolite, alpha-ketoglutarate (α- KG), showed a significant positive correlation with CLS. Although initial analysis also showed a significant correlation with RLS (p ≤ 0.02), it did not reach the significance threshold after p-value adjustment (Padj ≥ 0.05) (**Fig. 4A**). OA is an intermediate of *de novo* pyrimidine biosynthesis, which involves a series of reactions converting glutamine into ribonucleotides including UDP, CTP, thymine, and cytidine [51]. α-KG is a major TCA cycle intermediate, also involved in glutamic acid (Glu) and glutamine (Gln) synthesis, which can then be utilized for orotate and the synthesis of other amino acids. OA supplementation has been shown to extend lifespan in worms [52] and α-KG supplementation has been shown to increase lifespan in the nematode worm [53], fruit fly [54], and mice [55] and has been proposed as a central metabolite to treat aging and age-related diseases in humans [56]. To date, there have been no study demonstrating the longevity effect of OA and α-KG on yeast lifespan.

**Figure 4:**
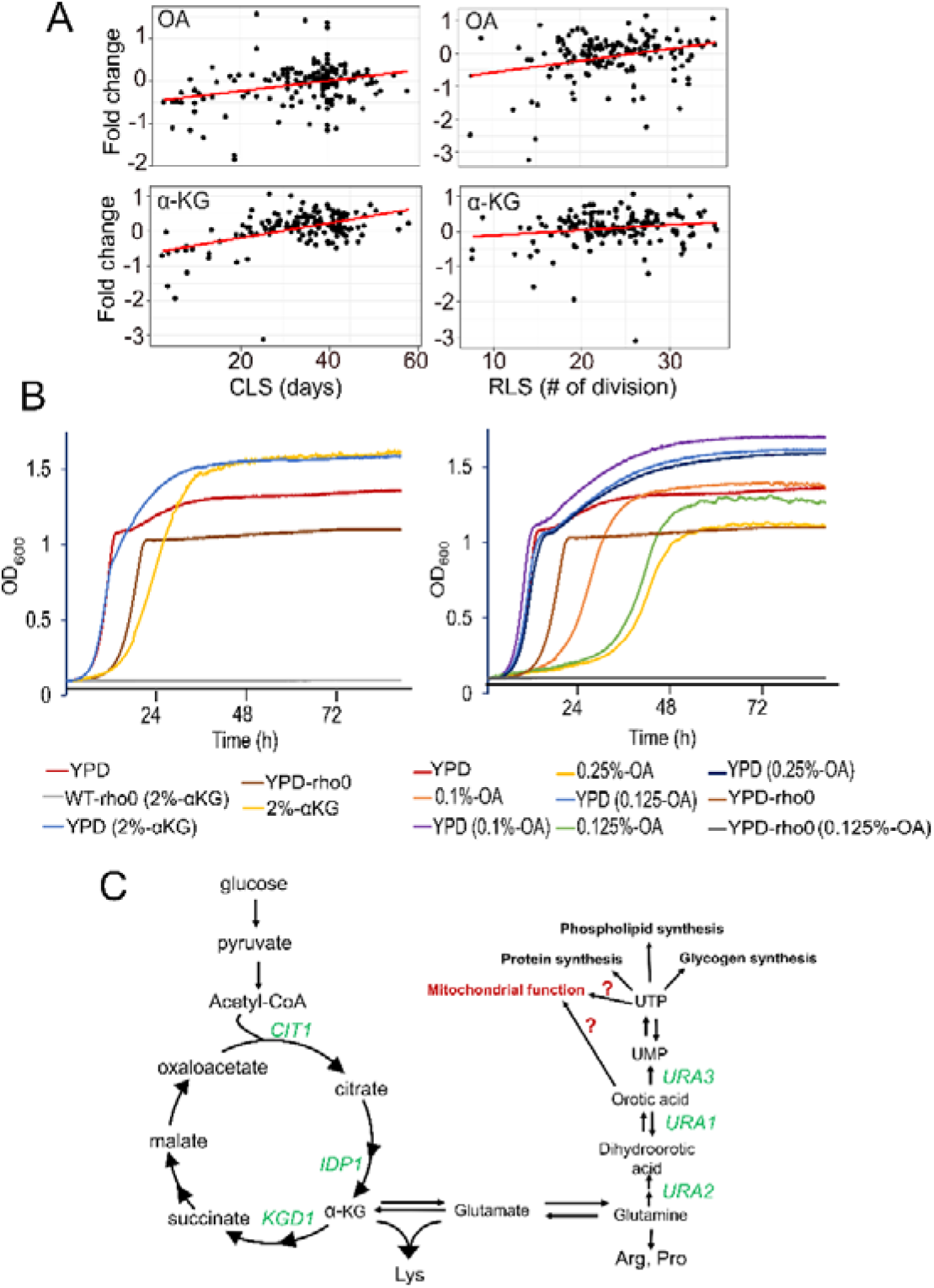
Association of Orotic acid (OA) and alpha-ketoglutarate (α-KG) abundances with lifespan and their effect on cell growth. (**A**) OA and α-KG abundances positively correlate with CLS (left) and RLS (right). P values and raw data can be found in Supplementary File 1. (**B**) Growth properties of WT strain in medium supplemented with α- KG (left panel) or OA (right panel) and effect of mitochondrial DNA (mtDNA) elimination (rho^0^) on α-KG and OA utilization. YPD indicates yeast peptone dextrose (2% glucose) medium. In the α-KG medium, glucose is replaced by 2% α-KG. Similarly, the OA medium replaces glucose with 0.1%, 0.125% or 0.25% OA. Each growth curve represents the mean values of five biological replicates (**C**) α-KG, and OA metabolism is depicted. Our data indicates both α-KG, and OA increase mitochondrial respiration. However, α-KG and OA-mediated mechanisms of increased mitochondrial function is currently not known (red arrows).

We first analyzed the growth phenotype of WT yeast on a medium supplemented with OA or α-KG. (**Fig. 4B**). We previously showed that wild budding yeast isolates could utilize α-KG as a carbon source, and it was associated with increased mitochondrial respiration in WT yeast isolates. However, after eliminating mtDNA completely, the yeast cells were no longer able to utilize α-KG [18]. To test whether a laboratory-adapted WT strain could also use OA and α-KG as a carbon source, we replaced glucose with 2% α-KG and 0.1%, 0.125% and 0.25% OA. To our surprise, we found that both α-KG and OA supplemented WT yeast grew in the absence of glucose (**Fig. 4B**). Supplementing the YPD medium (contains 2% glucose) with different concentrations of OA and α-KG improved growth phenotype in the stationary phase, further indicating these metabolites can be used as a carbon source in the absence of glucose. To further determine whether this effect was dependent on functional mitochondria, we eliminated mtDNA (rho^0^) in WT cells and repeated the growth analysis again. Our analysis showed that functional mitochondria were required for supporting growth in a medium supplemented with OA or α-KG (**Fig. 4B**). Overall, our analysis characterized these two metabolites as a respiratory carbon source that yeast can utilize to support its growth.

Next, to test whether the longevity effect of OA and α-KG is also conserved in yeast, we tested the RLS of WT yeast cells on a medium supplemented with OA or α-KG. Our result revealed that 2% α-KG supplementation in YPD did not cause significant lifespan alteration. However, eliminating glucose in the same media (see methods) significantly (Wilcoxon rank sum tests, p = 0.0001) increased median RLS by 29% (YPD: 26 versus α- KG: 34.5) (**Fig. 5A**). This data suggests α-KG alone can better support the full life cycle of WT strain and glucose abrogates the lifespan extension by α-KG in yeast. Similarly, we tested the OA effect on the lifespan of WT strain on a YPD medium supplemented with 0.1% OA or the same medium lacking glucose. We found that OA alone as a carbon source cannot support the full replicative life cycle of WT strain. On average, cells replicated 11-times (mean RLS). However, supplementing the YPD medium with 0.1% OA significantly (p = 0.006) increased median RLS by 17% (YPD: 27 versus YPD-OA: 31.5) (**Fig. 5B**).

**Figure 5:**
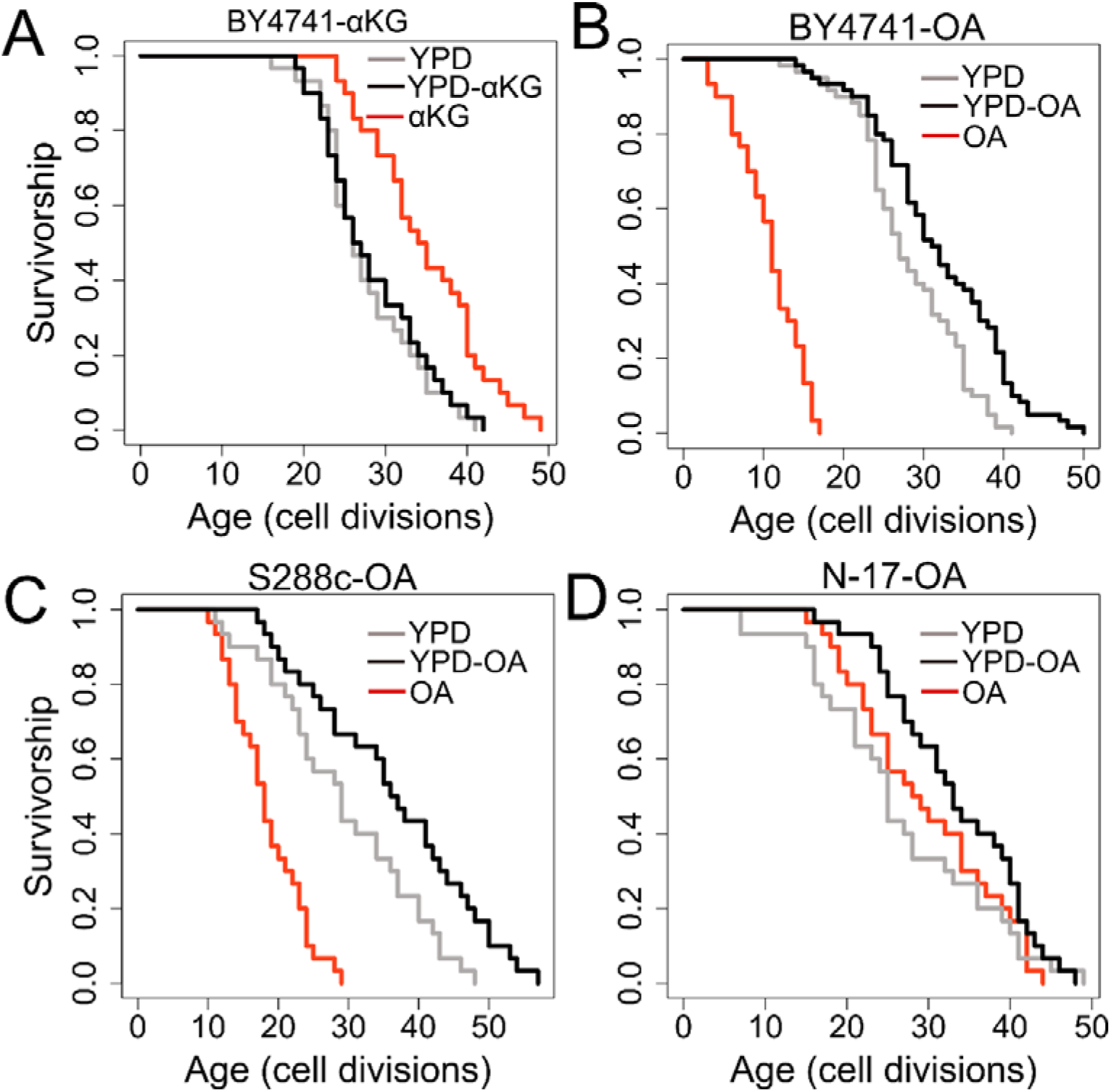
RLS effect of α–ketoglutarate and orotic acid supplementation. Lifespan curves for cells on medium supplemented with (**A**) α-KG or (**B, C, D**) OA. Standard YPD media contains 2% glucose. YPD-αKG and YPD-OA mediums were prepared by supplementing YPD with 2% αKG or 0.1% OA respectively. αKG and OA mediums contain 2% αKG or 0.1% OA as a carbon source. Lifespan data and the significance of lifespan changes can be found in Supplementary File 3.

The BY4741 strain is the derivate of S288c and harbors auxotrophic markers [57]. Among them is the *URA3* (Orotidine-5’-phosphate decarboxylase) gene, which catalyzes the sixth enzymatic step in the *de novo* biosynthesis of pyrimidines (**Fig. 4C**). To understand whether the OA effect on lifespan extension is an artifact of *URA3* auxotrophy, we also tested the lifespan of an original isolate of S288c strain under OA condition. We found that S288c cells can better utilize OA as a carbon in comparison to the BY4741 cells. On average S288c cells replicated 19-times (mean RLS) under OA condition and 29 times under glucose (YPD) condition. Adding 0.1% OA to YPD significantly (p = 0.03) increased median RLS. (YPD: 29 versus YPD-OA = 36.5) (**Fig. 5C)**. We further analyzed wild isolate of another budding yeast species, *Saccharomyces paradoxus*. Interestingly, our data showed that OA alone is sufficient to recapitulate the full replicative lifespan of the *S. paradoxus* strain (**Fig 5D**). Although not significant (p = 0.2), cells analyzed on 0.1% OA medium increased the median RLS by 14% (YPD: 25 versus OA = 28.5. Combining YPD with 0.1% OA resulted in significant (p = 0.01) median RLS increase (YPD: 25 versus YPD-OA: 33) (**Fig. 5D**). Overall, our data indicate that genetic background might be an important factor for the efficient utilization of OA as a carbon source. However, the actual mechanisms of OA utilization and the role of OA on mitochondrial function needs further studies.

## Discussion

Over the last three decades, both CLS and RLS models of yeast aging have contributed to the discovery of conserved longevity factors that modulate aging in higher eukaryotes [58, 59], but they are experimentally quite different paradigms that appear to be genetically regulated by largely distinct pathways [60, 61]. While it has been speculated that RLS may better reflect aging of mitotic tissues in higher eukaryotes and CLS better reflect post-mitotic tissue aging, RLS shares a greater degree of conservation with aging in the postmitotic nematode worm *C. elegans* compared to CLS [60, 62]. CLS and RLS are interconnected, however, as chronologically aged cells have a reduced RLS. Interestingly, the degree of reduction in RLS among chronologically aged cells at the single-cell levels is inversely correlated with mitochondrial membrane potential [63]. Mitochondrial function has also been shown to be critically important for normal CLS and RLS with several different mechanisms having been proposed, including mitohormesis, activation of mitophagy, mitochondrial retrograde signaling, and increased mitochondrial respiration [59, 64].

To understand how genetic perturbation of mitochondrial function affects aging, we assayed the chronological and replicative lifespan effect of single gene deletions corresponding to nuclear-encoded mitochondrial proteins. We found that, in general, mitochondrial genes whose knockout causes increased respiration, decreased translation, or amino acid biosynthesis, increase CLS, whereas those genes whose inhibition increase both cytosolic and mitochondrial ribosomes along with reduced actin depolymerization increase RLS. Interestingly, the CLS signature of mtKO strains resembles the signature of caloric restriction (CR), which has also been shown to increase mitochondrial respiration across different model organisms [17, 18, 20–23]. It should be noted, however, that increased CLS resulting from increased respiration is likely due primarily to reduced acidification of the culture medium during the fermentative growth phase. Future studies will be needed to establish whether such mechanisms are conserved outside of the yeast CLS paradigm [47].

The role of respiration in RLS and the response to CR in this context remains an open question. It has been proposed that CR increases RLS by activating the respiratory response and thereby activating Sir2 [65, 66]; however, other studies have shown that neither Sir2 nor respiration are required to achieve full lifespan extension from CR [67, 68]. Indeed, among a screen of single gene deletion mutants, several of the strains showing the largest RLS extension from CR corresponded to mitochondrial proteins [42]. Regardless of whether increased respiration is sufficient to increase RLS per se, our data indicate the plasticity of mitochondrial respiration can be targeted to regulate lifespan. For CLS, in particular, our data are well-aligned with previous findings that yeast CLS heavily depends on mitochondrial functional status [69].

One unexpected observation was that decreased activity of proteasome-dependent, ubiquitin-independent protein degradation positively correlates with CLS. This is in contrast with RLS, where activation of proteasome activity promotes longevity [70]. Most protein substrates are generally marked with polyubiquitin chains for degradation by the proteasome complex. However, denatured or partially unfolded proteins do not require ubiquitination for their degradation by the proteasome [71]. The data indicate that decreased activity of targeted protein degradation for such proteins might be beneficial for CLS. Furthermore, since the proteasome system is ATP-dependent [72], reduced proteasome activity might enhance intracellular ATP homeostasis in cells under stationary growth conditions.

We also observed significant variation in RLS across mtKO strains. We found that RLS y is strongly associated with up-regulation of oxidative phosphorylation, Fe-S cluster biogenesis, and ribosome biogenesis, including tRNA aminoacylation for protein translation and amino acid biosynthesis. Down-regulated pathways include actin filament depolymerization, endosome, trehalose metabolic process, and stress response. Previously, it has been shown that both cytosolic protein translation and actin dynamics are controlled by mitochondrial efficiency. In addition, mild inhibition of mitochondrial ribosomes was found to reduce cytosolic translation and modulates cytoplasmic protein quality control systems that determine cellular lifespan in yeast, worms, and mice [73, 74].

Lifespan extension upon a decrease in overall protein biosynthesis is a phenomenon that is conserved in different model organisms [75–77]. Contradictorily, we found that proteins involved in ribosome biosynthesis and components of cytosolic ribosomes involved in cytosolic translation are positively correlated with RLS, indicating increased but not decreased cytosolic translation machinery in long-living mtKO strains. Complete inhibition of nuclear-encoded mitochondrial proteins might activate mitochondria-derived mechanisms that attenuate both mitochondrial and cytosolic translation machinery, perhaps contributing to the loss of mitochondrial proteins. Recently, it has been shown that mitochondrial and cytosolic ribosomal proteins are balanced in a natural ratio, and a decrease in mitochondrial ribosomes in worms and mice reduces cytosolic ribosome abundances and translation [74]. In addition, mitochondrial translation efficiency was shown to be coupled with cytoplasmic protein homeostasis [73]. Our data also suggest that mitochondrial and cytosolic ribosome stoichiometry can still be regulated with increased abundances of ribosomes from both compartments without increasing global translation efficiency in the cytoplasm. In addition, increased ribosome density and increased tRNA biosynthesis might increase translation fidelity through tRNA selection during elongation and increase precise translation termination to prevent read-through [78]. This might decrease wasteful translational products (misfolded proteins) in long-lived mtKO strains. Along with the decreased stress response activity, activating mitochondrial translation might induce lifespan extension without reduced protein translation, and future studies are needed to uncover complete mechanisms. A similar observation was recently reported in nematode worms where a large-scale mutagenesis screen for increased survival revealed lifespan extension through integrated stress response inhibition without reduced mRNA translation [79]. Moreover, we recently showed that this observation of altered cytosolic translation and increased mitochondrial respiration is a general trend in long-lived laboratory-adapted yeast KO strains [18].

Finally, our analysis revealed that supplementation of two mitochondria-related metabolites, OA and α-KG, extend RLS. OA is an essential intermediate of pyrimidine *de novo* synthesis from glutamine (Gln). It has been shown that 0.5 mM OA supplementation induces a 15% lifespan extension in *C. elegans* by inhibiting reproductive signals and subsequently inducing the function of DAF-12 and DAF-16 [52]. The effect of OA supplementation on lifespan has not been tested in other organisms. In humans, dietary supplementation of OA has been suggested to provide a wide range of beneficial effects, including cardio protection and exercise adaptation [80].

Another longevity-promoting central metabolite, α-KG, has been shown to extend health span and lifespan in the nematode worm, fruit flies, and mice. This longevity effect was found to be mTOR-dependent in worms and fruit flies. However, whether the longevity effect of α-KG is mediated by TOR in mice needs further investigation [53–55]. To date, no study has analyzed the effect of α-KG on yeast lifespan. We previously showed that wild yeast isolates could metabolize α-KG as an alternative carbon source, and the elimination of mtDNA abolished their growth ability in the medium supplemented with α-KG. In addition, α-KG as a respiratory carbon source increased mitochondrial respiration [18]. Here, we found that increased α-KG abundance is associated with increased lifespan in mtKO strains. We further tested this observation by replacing glucose with α-KG in the yeast medium and analyzed RLS. In addition, we showed that utilization of both OA and α-KG requires functional mitochondria. This observation suggests a possibility that both metabolites might show their longevity effect by altering metabolic shifts from glycolysis to respiration. Furthermore, α-KG supplementation was shown to activate antioxidant defense, thereby preventing oxidative protein damage in yeast cells [81]. Additionally, *de novo* pyrimidine nucleotide synthesis might induce mitochondrial gene expression, as shown in mammalian cells [82], which, in part, might cause lifespan extension. However, the absence of the *URA3* gene in the BY4741 background suggests that the lifespan effect of OA cannot be attributed to increased pyrimidine nucleotide synthesis. This brings the possibility that increased mitochondrial function is the main reason for the extended lifespan. However, the mechanism that allows yeast cells to utilize OA as a respiratory carbon source and the downstream pathways involved in lifespan extension need further investigation.

One limitation of the experiments with α-KG presented here is that it entail simultaneous removal of glucose from the culture media. Reducing glucose levels from 2% to between 0.5 and 0.005% is sufficient to increase lifespan [66, 83]. Thus, we can not rule out the possibility that the observed lifespan extension results primarily from the absence of glucose, and α-KG is simply providing metabolic substrates to enable cell growth via non-fermentative processes. Future studies should be able to address this question by comparing various carbon sources for impact on yeast RLS and describing the genetic and biochemical mechanisms underlying the observed effects on lifespan. Our observation of background-dependent alternative carbon source utilization efficiency suggests that wild yeast isolates can be used as a discovery tool for deciphering the mechanisms.

Overall, we showed that the plasticity of mitochondrial function can be targeted for health and lifespan extension by genetic and metabolic interventions. Our data also indicates that regulatory pathways that mediate the longevity effect of OA and α-KG might be conserved among species and place-renewed emphasis on metabolic intermediates from mitochondrial pathways in aging biology. In addition, our data support that budding yeast may be used as a model organism further to dissect the longevity mechanisms of these two metabolites. Although, there has been much attention on beneficial effect of α-KG in humans [56, 84, 85], studies also showed that low dose of OA as a dietary component can exert beneficial properties such as cardiac rehabilitation, improved exercise tolerance in patients with coronary artery disease, neuroprotective role in brain injury and neurodegenerative disorders [51, 80, 86–88]. Our data from yeast lifespan analysis further suggesting the possibility that OA might have the potential to slow down aging in higher eukaryotes as well. Since both α-KG and OA are commonly used for muscle growth without any side effects, this brings the possibility of testing these metabolites in clinical trials for their impact on health parameters and known aging-associated pathologies.

## Materials and Methods

### Yeast strains and Media

Yeast strains lacking a single gene related to mitochondrial biology (mitochondrial knock-out strains-mtKO strains) (**Supplementary Table 1**) were obtained from the haploid yeast ORF deletion collections [89], and the WT (BY4741; *MAT**a** his3Δ1 leu2Δ0 met15Δ0 ura3Δ0*) was purchased from American Type Culture Collection (ATCC). Wild isolate of S288c and S. paradoxus strain, N-17 were previously described [18]. Cells were grown on standard YPD containing 1% yeast extract, 2% peptone, and 2% glucose at 30°C, or YPD supplemented with 2% αKG or 0.1% OA. Alternative carbon source plates were prepared by combining 2% peptone, 1% yeast extract, 2% agar and 2% αKG or 0.1% OA.

### Data Acquisition

The relative levels of proteins, metabolites, and lipids in mtKO strains under the two culture conditions (expressed as log2 fold-change with respect to wild type) had been previously quantified by mass spectrometry [35], so the raw data were downloaded from a publicly available database. After excluding those molecules with missing values in >20% of the strains and filling up the remaining missing values, we performed KNN imputation [90]. As a result, we obtained the datasets of 1931 proteins, 219 metabolites, and 53 lipids under respiratory condition and 1874 proteins, 183 metabolites, and 52 lipids under fermentation condition (**Supplementary Table 2**). The corresponding RLS data for mtKO strains were from the previously published data set [61]. Based on the raw data from the number of replicates, we calculated mean and maximum RLS, and the mean RLS values were used to identify genes and metabolites associated with RLS across mtKO strains.

### Yeast Chronological Lifespan

Outgrowth-based yeast CLS assays were performed as previously described elsewhere [91–93]. A BioTek EPOCH2 machine was used for all outgrowth assays. In short, single colonies for each mtKO strain were grown in 200 μl (96-well microplates) YPD medium overnight at 30°C. The next day aliquots of 2 μl of each culture were inoculated into new YPD medium and then subjected to 48-hour automated kinetic growth assay with OD_600_ monitoring with the shaking module set to high continuous shaking. After six weeks of growth analysis, doubling times and lag times were extracted from the generated growth curves and used to determine the proportion of living cells in the aging cultures for each mtKOs in biological triplicate using YODA software as previously described [91, 94].

### Correlation relationship between omics and lifespan phenotypes

To identify genes and molecules whose levels are correlated with lifespan phenotypes, we use Spearman’s rank-order correlation. Spearman’s correlation coefficient rho provides a nonparametric measure of the strength and direction of association between the two variables. We performed Spearman’s correlation between respiration omics and CLS, fermentation omics, and RLS and computed the adjusted p values by multiple testing correction using FDR for significance. Principal component analysis was performed on imputed values R package stats. Functional enrichment analysis of the significant gene lists from both conditions were done using gProfiler [95].

### Experimental Testing of Lifespan Extending Effect of α-KG and Orotate Supplementation

To investigate the lifespan effect of α-KG and orotate, we performed a RLS assay on solid mediums supplemented with each metabolite. RLS was determined using our previously published protocol [18] with some pre-conditioned modification for α-KG and OA. Briefly, yeast cell culture was freshly started from frozen stocks on YPD plates and grown for two days at 30 °C. Prior to dissection, several colonies were pre-cultured in liquid medium lacking glucose and supplemented with α-KG (2%), or orotate (0.1%) for three days. The next day 25 μl aliquots from each culture were dropped onto their matching agar plates and incubated two days. After then small amount of cells from each plate transferred to new aging plate with pipette tips and subjected to RLS analysis. Around ~100 dividing cells were lined up and after the first division, newborn daughter cells were chosen for RLS assays using a dissection microscope. Generation of survival curves and statistical analyses were performed using the survival and flexsurv packages in R, respectively. Finally, for generating rho^0^ cells, mtDNA was eliminated by culturing cells in YPD medium, supplemented with ethidium bromide [18].

